# Silver chimaera genome assembly and identification of the holocephalan sex chromosome sequence

**DOI:** 10.1101/2024.08.20.608884

**Authors:** Akinori Teramura, Mitsutaka Kadota, Shotaro Hirase, Shigehiro Kuraku, Kiyoshi Kikuchi

## Abstract

Cartilaginous fishes are divided into holocephalans and elasmobranchs, and comparative studies involving them are expected to elucidate how variable phenotypes and distinctive genomic properties were established in those ancient vertebrate lineages. To date, molecular-level studies on holocephalans have concentrated on the family Callorhinchidae, with a chromosome-scale genome assembly of *Callorhinchus milii* available. In this study, we focused on the most species-rich holocephalan family Chimaeridae and sequenced the genome of its member, silver chimaera (*Chimaera phantasma*). We report the first chromosome-scale genome assembly of the Chimaeridae, with high continuity and completeness, which exhibited a large intragenomic variation of chromosome lengths, which is correlated with intron size. This pattern is observed more widely in vertebrates and at least partly accounts for cross-species genome size variation. A male-female comparison identified a silver chimaera genomic scaffold with a double sequence depth for females, which we identify as an X chromosome fragment. This is the first DNA sequence-based evidence of a holocephalan sex chromosome, suggesting a male heterogametic sex determination system. This study, allowing the first chromosome-level comparison among holocephalan genomes, will trigger in-depth understanding of the genomic diversity among vertebrates as well as species’ population genetic structures based on the genome assembly of high completeness.

## Introduction

Chimaeras (subclass Holocephali) are classified into three families, namely Chimaeridae, Rhinochimaeridae, and Callorhinchidae; the subclass Holocephali together with the subclass Elasmobranchii comprise the vertebrate class Chondrichthyes (reviewed in Finucci et al., 2021). Investigations of holocephalan species are expected to provide valuable information about the evolutionary origins of these deep-sea organisms and their ecological interactions as well as the evolution of vertebrate genome. To date, however, genomic investigations on this taxon have been limited to a single species, the elephant fish (*Callorhinchus milii*; often called as elephant shark) (Venkatesh et al., 2014).

*Chimaera phantasma* (Family Chimeridae; Figure 1) is a demersal deep-sea chimaera that lives in the North Pacific. Since most species in this family are deep-sea dwellers and are thus highly elusive, it is not trivial to secure tissue samples for extraction of high-quality DNA and intact RNA. However, in some parts of Japan, *Ch. phantasma* dwells in relatively shallow waters, approximately 100 meters in depth, and is more easily obtained. This species is listed as “Vulnerable (VU)” on the IUCN Red List, because it has been strongly affected by human activities despite the lack of biological studies (reviewed in Finucci et al., 2021). Demography of this species as well as its population structure can be assessed more reliably using high-quality genome assembly.

**FIGURE 1.**
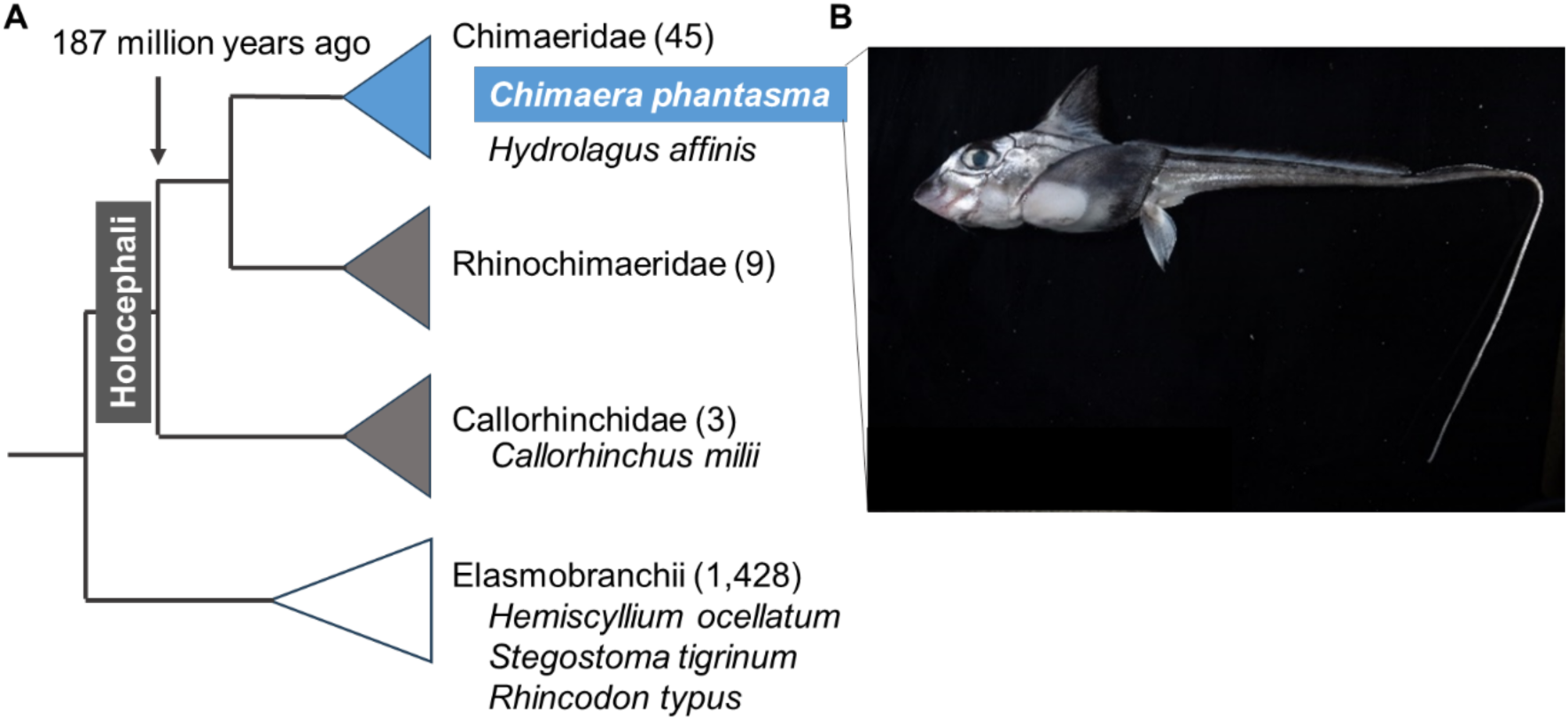
*Chimaera phantasma*. A, Schematic phylogeny. Parentheses include the numbers of described species as of January 2024 (Fricke et al., 2023). B, Adult individual of *Ch. phantasma*. Photo credit: Akinori Teramura.

*Chimaera phantasma* and *Ca. milii* are estimated to have diverged approximately 187 million years ago (Inoue et al., 2010). Cytogenetic analyses identified highly variable karyotypes within Chimaeriformes: the spotted ratfish (*Hydrolagus colliei* ) has a diploid chromosome number of 58, while the rabbit fish (*Chimaera monstrosa*) has a diploid chromosome number of 86 (Stingo and Rocco 2001).

Genome sizes of holocephalan species are smaller than those of other cartilaginous fishes (Venkatesh et al., 2005), in contrast to chondrichthyan species such as sharks and rays whose genome sizes often exceed 3 Gb (reviewed in Kuraku, 2021). *Callorhinchus milii* was used for whole genome sequencing (Venkatesh et al., 2007) and was the only holocephalan species whose whole genome had been sequenced (Venkatesh et al., 2014) until that of the rabbitfish (*Hydrolagus affinis*) became available in 2020 (GCA_012026655.1). As the *H. affinis* genome assembly remains at the contig-level (Table 1), chromosome-level genome sequence information of Holocephali is still limited to *Ca. milii* (Nakatani et al., 2021). Genomic resources for more holocephalan species will permit comparative investigations on the variation of genome sizes and karyotypes as well as possible polyploidization. Moreover, it is of great interest to relate DNA-level variations with variations in habitat. Another aspect of holocephalan genomics that requires study is that of sex determination. In 2023, the first chondrichthyan sex chromosome was reported from zebra shark (*Stegostoma tigrinum*) and whale shark (*Rhincodon typus*); in both species, the X chromosome was identified (Yamaguchi et al., 2023). This report was followed by the Y chromosome identification for white-spotted bamboo shark (*Chiloscyllium plagiosum*) (Wu et al., 2024). On the other hand, sex chromosomes of holocephalans have never been documented. It is likely that holocephalans have a different mode of sex chromosome organization from elasmobranchs after 400 million years of divergence.

**TABLE 1.** Basic statistics of *Chimaera phantasma* genome assemblies and comparison with other holocephalan genome assemblies.

| Species | *Chimaera phantasma* | *Callorhinchus milii* | *Hydrolagus affinis* |
| --- | --- | --- | --- |
| Assembly ID<br>(Accession ID) | sChiPha1.1 | IMCB_Cmil_1.0<br>(GCF_018977255.1) | UP_Haf<br>(GCA_012026655.1) |
| Sequencing technology | PacBio<br>CLR* | PacBio<br>CLR | Illumina<br>short reads |
| Assembler | wtdbg2 | Falcon | W2RAP v. 1.0 |
| Proximity ligation method | iconHi-C | Dovetail Chicago,<br>Dovetail Hi-C | Not used |
| Scaffolder | 3d-dna | HiRise | Not used |
| Total number of bases (Mb) | 1,010 | 941 | 1,114 |
| Single copy ortholog<br>completeness<sup>#</sup> | 96.8 | 93.1 | 56.4 |
| Max. scaffold length (Mb) | 155.4 | 139.2 | 0.397 |
| N50 scaffold length (Mb) | 42.7 | 69.3 | 0.019 |
| % of bases in scaffolds > 1 Mb | 96 | 90.8 | 0 |
| Number of sequences > 1 Mb | 38 | 31 | 0 |
| % Gaps | 0.04 | 0.01 | 0 |

<sup>#</sup>Completeness was measured with the program pipeline compleasm (Huang & Li, 2023).
\*CLR, continuous long read.

In this study, we sequenced the whole genome DNA of the silver chimaera (*C. phantasma*). We used it to consider the process of genome evolution in the two chondrichthyan lineages, namely Holocephali and Elasmobranchii. We show that their genomes have maintained high repetitiveness and within-karyotype chromosome-size variation, while they exhibit a remarkable difference in genome size, which is partly contributed by intron length variation. We also report the first genetic evidence of male heterogamety in a holocephalan species as we identified a 500-kb sequence with a double sequence depth for females, which was revealed by exhaustive genome resequencing of multiple individuals of both sexes. Comparisons of the holocephalan sex chromosome with the sequences reported for elasmobranch species as well as those of mammals and birds suggest the independent origin of sex chromosomes in these lineages.

## Methods

### Genome sequencing and assembly

A young female *Ch. phantasma* caught in Suruga Bay, Shizuoka prefecture as a commercial trawl bycatch was used to obtain high molecular weight DNA using the phenol/chloroform method with TNES-urea buffer (Asahida et al., 1996). The concentration of the extracted DNA was measured with a Qubit Fluorometer (Thermo Fisher Scientific, MA, USA), and the size distribution of the DNA fragments was analyzed with TapeStation 2100 (Agilent Technologies, CA USA) to ensure high integrity. An SMRT sequence library was constructed with an SMRTbell Express Template Prep Kit 2.0 (Pacific Biosciences, CA, USA) and was sequenced in a single 8M SMRT cell on a PacBio Sequel II system (Pacific Biosciences) at Macrogen Inc. A total of 77.7 Gb continuous long reads (CLR) were assembled using the program wtdbg v2.5 with its default parameters (Ruan & Li, 2020).

### Hi-C data production and genome scaffolding

An Hi-C library was prepared using muscle tissue from the female (named sChiPha1) used for genome sequencing as described in the the iconHi-C protocol (Kadota et al., 2020). The library was digested with the restriction enzymes DpnII and HinfI, and was sequenced on a HiSeq X sequencing platform (Illumina Inc., CA, USA). This sequencing produced 263 million PE150 read pairs that amounted to 79 Gb in total. The obtained Hi-C read pairs were processed with the program Trim Galore! v0.6.8 (https://www.bioinformatics.babraham.ac.uk/projects/trim_galore/) specifying the options ‘--phred33 --stringency 2 --quality 30 --length 25 --paired’ and aligned to the HiFi sequence contigs with the program Juicer v1.6. HiFi sequence contigs were scaffolded with 3d-dna (version 201008; Dudchenko et al., 2017) specifying the options ‘-m haploid -i 1000 --editor-repeat-coverage 15 -r 2’ to be consistent with the chromatin contact profiles. The continuity and completeness of the resultant genome assembly, designated as sChiPha1.1, as well as the predicted protein-coding gene set, was assessed with the webserver gVolante v2.0.0 (Nishimura et al., 2017) in which the pipeline BUSCO v5.1.2 was implemented (Seppey, Manni, and Zdobnov 2019), using the ortholog set ‘vertebrate_odb10’ supplied with BUSCO. The completeness of the nucleotide sequences of the genome assemblies was assessed with compleasm (Huang & Li, 2023) (formerly called minibusco) that is designed to achieve higher accuracy through use of miniprot (Li 2023).

## Repeat identification

The obtained *Ch. phantasma* genome assembly was subjected to *de novo* repeat element identification with RepeatModeler ver. 2.0.4 (Flynn et al., 2020) with the option ‘-LTRStruct’, and the resultant repeat library was input into repeat masking using RepeatMasker ver. 4.1.5 with the default parameters (Smit et al., 2013).

### Transcriptome sequencing

Total RNA (0.5–2.7 μg) was extracted using an RNeasy Plus Universal Mini Kit (Qiagen, Hilden, Germany) from brain, kidney, ovary, gill, liver, heart, and muscle from a second adult female sampled from the East China Sea, off Nagasaki prefecture. RNA-seq was performed on total RNAs from each tissue. Libraries were prepared using the TruSeq stranded mRNA Library Prep Kit (Illumina, CA, USA) and sequenced on NovaSeq 6000 to obtain PE reads with 150 cycles at Macrogen Inc.

### Comparative analysis of the protein-coding gene landscape

The RNA-seq reads obtained above were mapped to the *Ch. phantasma* genome assembly using the program STAR Version 2.5.2b (Dobin et al. 2013), and the resultant bam file was input as evidence of transcription in protein-coding gene prediction to Braker version 2.1.5 (Hoff et al. 2015) incorporating all peptide sequences in the file, odb11_vertebrata_fasta, provided by OrthoDB (Kuznetsov et al. 2023).

Structural assessment of protein-coding gene structures and their genomic distributions were demonstrated using the general feature format (GFF) file from the above-mentioned Braker2 analysis as well as GFF files retrieved from NCBI Genomes for the following species: *Ca. milii* (Accession ID, GCF_018977255.1), epaulette shark *Hemiscyllium ocellatum* (GCA_020745735.1), Japanese medaka (GCF_002234675.1), chicken (GCA_016699485.1), and human (GCF_009914755.1). After information for intronic regions were amended by the program agat_sp_add_introns.pl (Dainat et al., 2023) for the epaulette shark, the lengths of individual introns were analyzed from these GFF files to compute length distributions for individual chromosome sequences for each species.

### Orthology inference and cross-species comparisons of chromosomes

The above-mentioned protein-coding gene collection for the six species were modified to retain only peptide sequences with the longest open reading frames for individual genes; these were used as input for orthogroup construction with Orthofinder v2.5.5 (Emms and Kelly, 2019). Homology of chromosomes between species was inferred using the locations of candidate orthologous genes classified by Orthofinder.

### Whole genome resequencing for sex chromosome identification

Genomic DNAs were extracted from nine males and nine females from East China Sea, off Okinawa prefecture to off Chiba prefecture and used to obtain paired-end short reads of 20–30x fold with PE150 on the DNB-seq sequencing platform at Macrogen Inc. (Supplementary Table 1). The total lengths of sequenced reads for females were 28.3–29.0 Gb and for males were 20.0–26.0 Gb. These reads were mapped to the genome assembly sChiPha1.1 with the mem option of bwa version 0.7.17-r1188 (Li & Durin, 2009), and the sequencing coverages from these BAM files were quantified with the bamcoverage option of deeptools version 3.5.4. (Ramírez et al., 2016). The average coverage in 10-Kb windows was compared between females and males.

## Results

### Genome sequencing and assembly

Continuous long-read (CLR) sequencing of the genome of silver chimaera (Ch. phantasma) produced a total read size of 77.7 Gb (76.9x of the whole genome if the genome size is assumed to be 1.01 Gb) with an average length of 12.9 Kb. These reads were assembled into 2,074 contigs with an N50 length of 11.4 Mb; 29 sequences exceeded 10 Mb, with a maximum sequence length of 54.6 Mb. We separately prepared an Hi-C library and obtained 78 Gb paired-end read data (263 M PE150 pairs) from this library. The Hi-C data were used in Hi-C scaffolding into a chromosome-scale genome assembly (sChiPha1.1) containing 2,094 sequences with an N50 length of 42.7 Mb (Figure 2) and a maximum sequence length of 155.4 Mb (Table 1). In comparison with other genome assemblies of chimaeroid species, our genome assembly exhibited a higher completeness of genomic regions in chromosome-scale sequences (proportion of total bases in the scaffolds longer than 1 Mb: 96.0% versus 90.2% for *Ca. milii*) and of conserved protein-coding genes shared as single-copy genes between vertebrates (completeness of single copy orthologs: 96.8% versus 93.1% for *Ca. milii*) (Table 1). The chromosomal organization in *Ch. phantasma* resembles that of *Ca. milii* in that a relatively small number of chromosomes stand out in length: four chromosomes longer than 85 Mb and about 30 chromosomes shorter than 45 Mb. This pattern confirms cytogenetic analyses on other chimaeroid species that reported diploid chromosome numbers of 58 and 86 (*H. colliei* and *Ch. monstrosa*, respectively) (Stingo and Rocco 2001). Despite our best efforts, the karyotype of *Ch. phantasma* has not yet been obtained. The genomes of *Ch. phantasma* and *Ca. milii* share a gradual variation in length of chromosomal sequences, which also resembles that of the chicken (Figure 3A).

**FIGURE 2.**
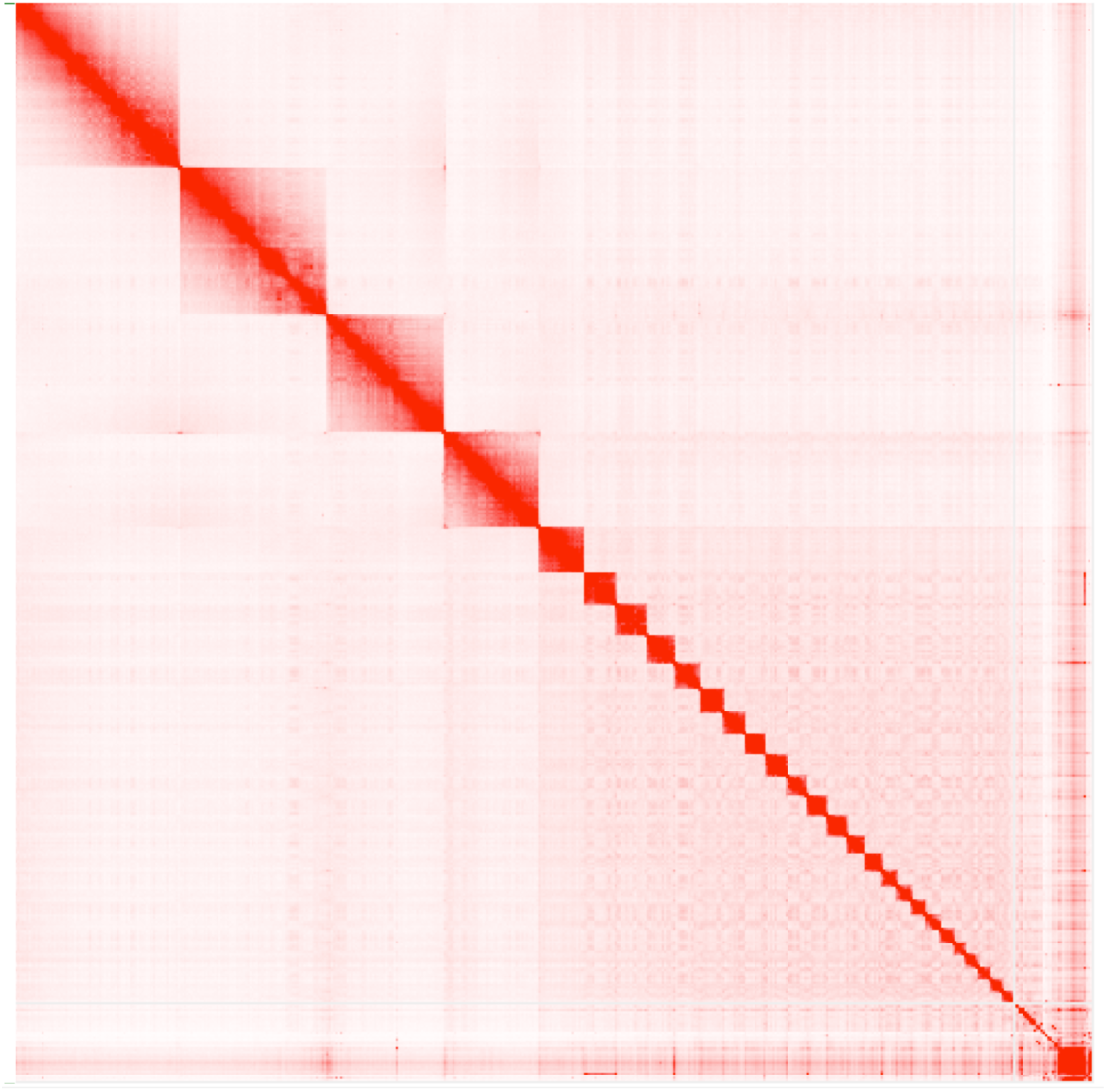
Hi-C contact map for *Ch. phantasma*. This genome consists of four chromosomes longer than 85 Mb and about 30 chromosomes shorter than 45 Mb.

**FIGURE 3.**
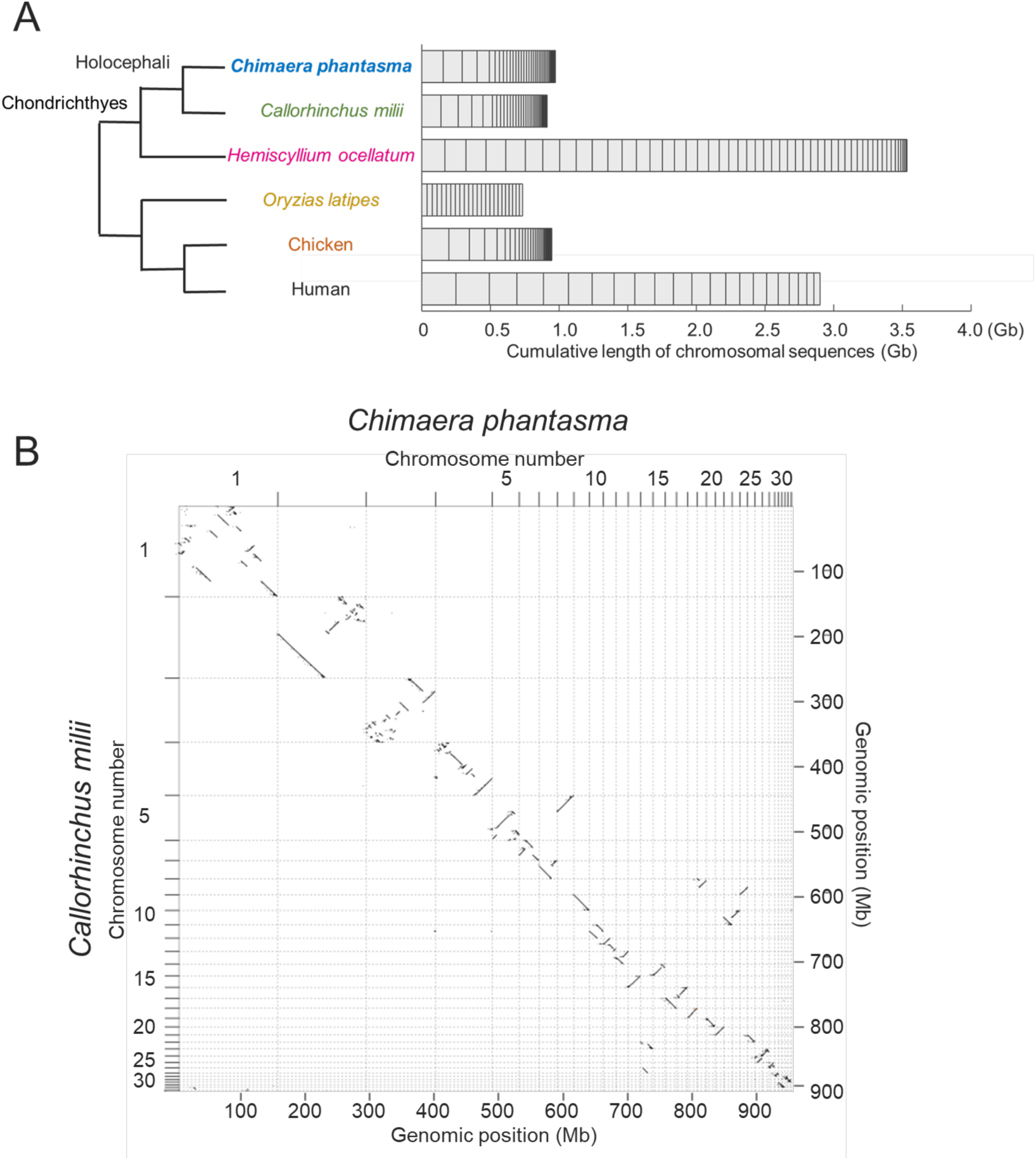
Comparative overview of genomic organization. A, Breakdown of whole genome assembly size into different chromosomal scaffolds. Chromosomal sequences are sorted by length and scaffolds shorter than 0.2 Mb are excluded. B, Cross-species comparison of whole genome organization between *Ch. phantasma* and *Ca. milii*. Sequences of high similarity are shown with diagonal lines by the program D-GENIES (Cabanettes & Klopp, 2018) with the “many repeats” mode.

We performed the first-ever cross-species comparison of chromosomal sequences within the Holocephali in a comparison of *Ch. phantasma* and *Ca. milii* (Figure 3B). This comparison was based on nucleotide sequence alignments and highlighted high homology of *Ch. phantasma* chromosome 1 and *Ca. milii* chromosome 1, despite the intra-chromosomal breaks (Figure 3B). Likewise, this comparison showed a one-to-one relationship for most chromosomes (except for chromosomes 5, 8, and 10 of *Ca. milii,* each of which corresponded to two *Ch. phantasma* chromosomes).

### Characterization of genomic compositions

A previous vertebrate-wide comparison revealed remarkable intragenomic heterogeneities in base composition and the distributions of protein-coding genes and repetitive elements (Yamaguchi et al., 2023). In this study, we performed a *de novo* repeat library construction and detection of repetitive elements in the *Ch. phantasma* genome assembly, and compared the repeat landscape with that of other chondrichthyan genomes. This analysis revealed a similar profile in the genomes of *Ch. phantasma* and *Ca. milii* (Figure 4). Although the total amounts of repeat elements (in both copy numbers and total bases) are much smaller in the two chimaera species than in elasmobranchs and are dependent on genome size, the overall proportion of repetitive elements in the whole genome is maintained throughout chondrichthyan species (Figure 4).

**FIGURE 4.**
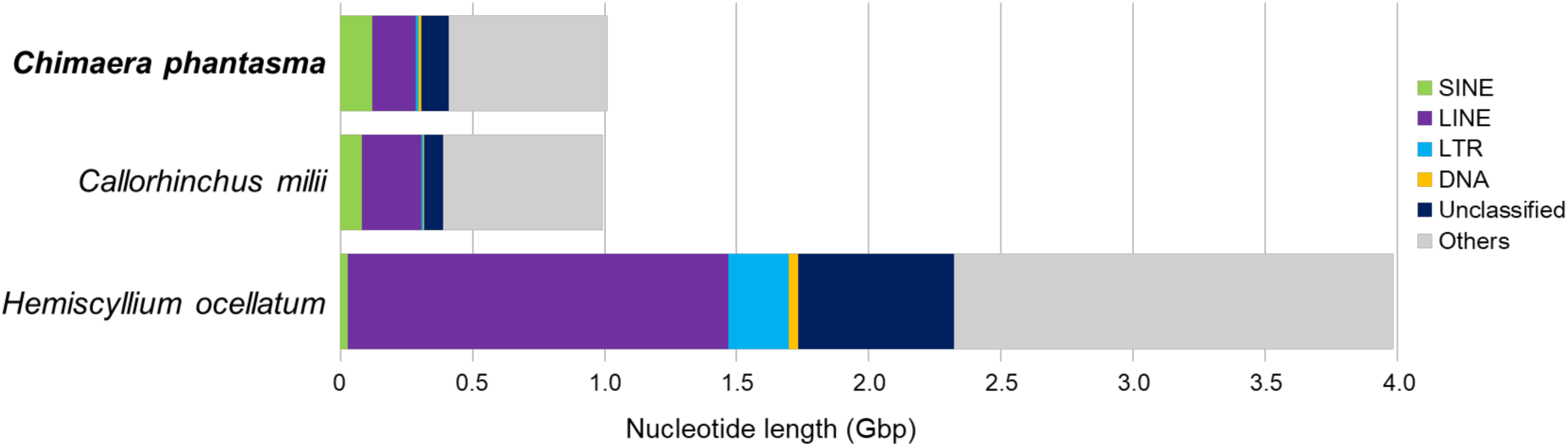
Comparison of the repetitive element breakdown in two chimaera and one shark species. Proportions of genomic regions annotated to different repeat classes were obtained using RepeatModeler and RepeatMasker as described in the Methods.

### Cross-species comparison of protein-coding landscape

In the *Ch. phantasma* genome assembly obtained here, exons for protein-coding genes were inferred by incorporating evidence of transcription from the RNA-seq data, as well as peptide sequences from other species (see Methods). In total, 24,724 protein-coding gene models were identified; this estimate resembles those obtained in other vertebrates.

To assess the effect of genome size variation on gene structure, we focused on intron lengths. A previous analysis of elasmobranch genomes showed they had larger introns compared to other vertebrates including *Ca. milii* (Hara et al., 2018). It will be of great interest to identify which genome sequences are particularly responsible for this intron enlargement. In other shark species, large chromosomes tend to harbor large genes (including both exons and introns) (Yamaguchi et al., 2023); however, no analysis to date has demonstrated a chromosome-length-dependent pattern with respect to introns. For this reason, we compared the median length of introns in the predicted protein-coding genes between different *Ch. phantasma* chromosomes. We found a clear tendency for longer chromosomes to harbor protein-coding genes with longer introns (Pearson’s correlation coefficient r = 0.96; Figure 5). This pattern is also observed in *Ca. milii*, *H. ocellatum*, and chicken, whereas human and medaka genomes do not exhibit this trend (r <0.25; Figure 5). This analysis suggests that the length variation of chromosomes can at least partly be linked with intron length variation, and its extent may partly account for the variation of whole genome sizes.

**FIGURE 5.**
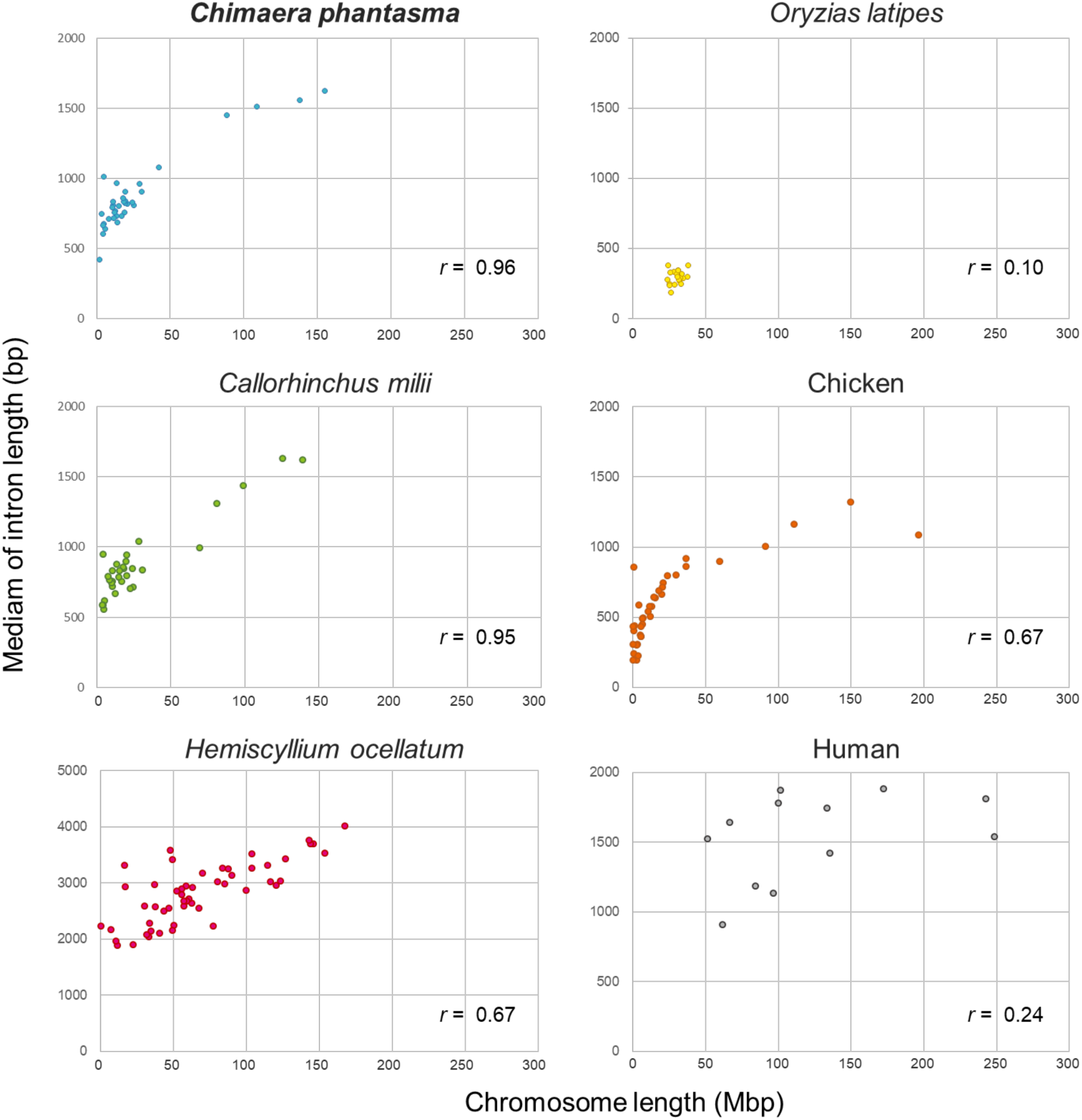
Intron size per chromosome. The vertical axis shows the median intron size and the horizontal axis shows the chromosome length. Sequences longer than 2 Mbp are regarded provisionally as chromosomes in the sChiPha1.1 assembly (see Discussion). Note that the scale for the vertical axis of *H. ocellatum* differs from that of the other panels. The Pearson’s correlation coefficients in the panels were calculated by the cor function of R ver. 4.0.3.

In the scatter plot shown in Figure 5, some outliers are present (Figure 5). These chromosome outliers in chondrichthyan species tend to harbor a small number of protein-coding genes: in *Ch. phantasma*, chromosome 35 (2.09 Mb) contains only three predicted protein-coding genes; in *Ca. milii* in which sequences shorter than 1 Mb are labelled as ‘chromosomes’ (Nakatani et al., 2021), chromosomes 34 (933 Kb) and 35 (792 Kb) contain only four and five genes, respectively. The paucity of genes on these ‘chromosomes’ prevents us from judging whether these sequences have unusual genomic characteristics or if this is caused by the small sampling size. It is also possible that these are fragments of longer chromosomes that were not assembled or scaffolded properly.

### Identification of sex chromosome sequences

Holocephalan sex chromosomes have not previously been identified in either cytogenetic or genomic analyses. This prompted us to search for candidate sex chromosomes in our *Ch. phantasma* genome assembly. For this purpose, we performed whole genome resequencing of eight male and female *Ch. phantasma* (that did not include the individual used for long-read sequencing), and quantified the sequencing coverages of individual genome scaffolds in the assembly sChiPha1.1. The comparison between the males and females revealed a ratio of approximately 0.5 for *Ch. phantasma* Scaffold 41 (Figure 6), suggesting this scaffold was sequenced twice in females compared to other genomic regions. Assuming this species has an XY sex chromosome system (male heterogamety), Scaffold 41, which is about 500 Kb-long, might represent a fragment of the X chromosome. Although we observed other genomic regions with deviating ratios, scaffold 41 was the only one with a consistent deviated pattern in its whole sequence stretch.

**FIGURE 6.**
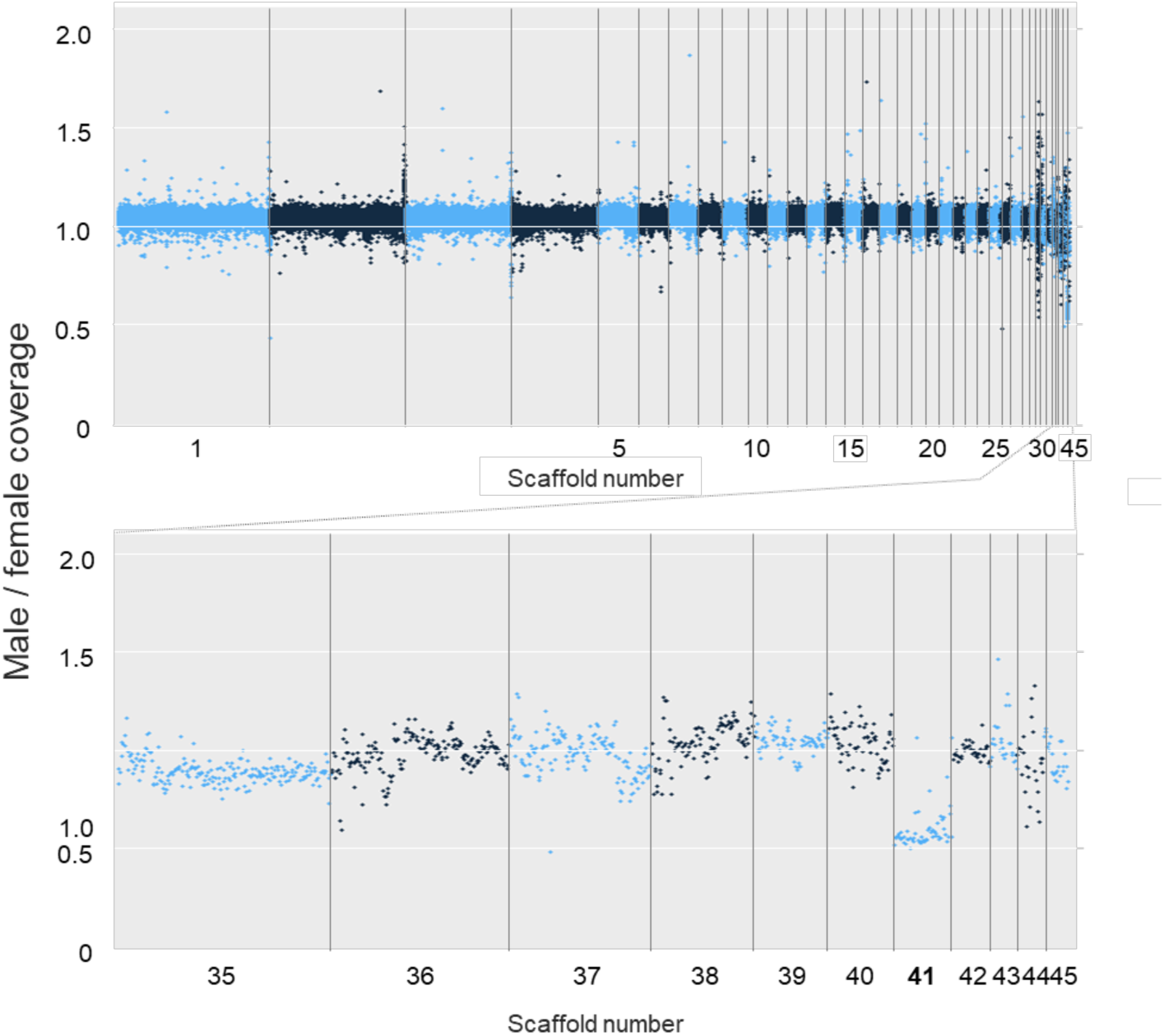
Male-female comparison of sequencing depth for identifying sex-related genomic regions. The upper panel shows all the scaffolds and the lower panel shows the part of them.

The *Ch. phantasma* Scaffold 41 is predicted to harbor 61 protein-coding genes. These genes do not include any orthologs of genes identified as master sex determination genes in other species (Graves, 2006; Bertho et al., 2021; Nagahama et al., 2021; Kitano et al., 2024). Orthologs of the 61 genes are located on chromosome 1 of the human genome and chromosome 2 of the chicken genome. Conversely, most orthologs of genes on the human X and Y chromosomes are located on Scaffolds 4 or 13 of the *Ch. phantasma* genome, and most of those on chicken Z and W chromosomes are found on Scaffold 1 of the *Ch. phantasma* genome. This comparison with human and chicken genomes shows the distinct autosomal origin of the *Ch. phantasma* X chromosome compared to those non-chondrichthyan lineages with established sex chromosomes.

Many of the orthologs of the genes predicted on *Ch. phantasma* scaffold 41 are located on chromosome 50 of the *H. ocellatum* genome and chromosome 47 of the *S. tigrinum* genome; these two chromosomes are not sex chromosomes in these elasmobranch species. The *S. tigrinum* X chromosome has been shown to be homologous to human chromosome 12 and to chicken chromosome 34 (Yamaguchi et al., 2023). This putative elasmobranch X chromosome harbors *Dhh*, *Gdf11*, a cluster of *Wnt1*, *-6b*, *-10b* genes in its X-specific region, as well as HoxC genes in its pseudoautosomal region. A search for a region in the *Ch. phantasma* genome that is homologous to this *S. tigrinum* X chromosome consistently identified *Ch. phantasma* Scaffold 29 (6.1 Mb). This scaffold harbors orthologs of *Dhh*, *Gdf11*, *Wnt1*, *-6b*, -*10b*, and *HoxC*. The *Ca. milii* genome possesses a highly constrained Hox C cluster as in many other vertebrates (Ravi et al., 2009; reviewed in Kuraku 2023); we found that the *Ch. phantasma* genome likewise harbors a conserved Hox C cluster with identical gene repertoires (*HoxC1*, -*C3*, -*C4*, -*C6*, -*C8*, -*C9*, -*C10*, -*C11*, -*C12*, and -*C13*).

## Discussion

### Trends in holocephalan karyotypes from a DNA-based viewpoint

The subclass Holocephali is species-poor with a maximum of 60 species; however, it occupies a unique phylogenetic position with approximately 400 million years of divergence from the Elasmobranchii, the other extant chondrichthyan lineage. In concordance with the results from cytogenetic studies on other holocephalan species (Stingo and Rocco 2001), we found that the *Ch. phantasma* genome assembly contained only four chromosomal sequences longer than 85 Mb, while the fifth longest sequence was shorter than 45 Mb. A relatively large gap between the sizes of the large and small chromosomes may be a characteristic of holocephalan species, given that the genome assembly of *Ca. milii* also showed a similar pattern of chromosome size variation (Nakatani et al. 2021). However, this interpretation needs to be supported by information from other species, especially those in the Rhinochimaeridae for which no genome assembly (even on a non-chromosome level) was available as of July 2024.

### Genomic organization within chondrichthyans

In comparison with those of other vertebrates, chondrichthyan genomes have a wider distribution of chromosome lengths, with dozens of chromosomes being shorter than 30 Mb (Figure 3). A high number of short chromosomes in karyotypes has also been observed in most bird and reptile species; these small chromosomes are termed ‘microchromosomes’. This pattern of karyotype variation is not seen in teleost fish or mammalian species. Nevertheless, there are remarkable differences among chondrichthyan species: the analyzed holocephalan species have only a few chromosomes longer than 50Mb, while elasmobranchs have dozens (Yamaguchi et al., 2023; Sendell-Price et al., 2023; Marletaz et al., 2023).

The range of genome sizes differs between Holocephali and Elasmobranchii. Species in the latter taxon often have genome sizes larger than 3 Gb, with a current maximum of 15 Gb (reviewed in Kuraku, 2021), while the genomes of holocephalans are reported to be smaller (Venkatesh et al., 2005). This is exemplified by the relatively small size (1.01 Gb) of the *Ch. phantasma* genome assembly produced here, and by that of *Ca. milii* (Table 1). The one caveat to this conclusion is that the number of holocephalan genomes that have been investigated remains very small. A Feulgen staining analysis of the genome size of *Ca. milii* provided an estimate of 1.9 Gb (Hardie and Hebert, 2004). This is clearly larger than the genome assembly size (Table 1). Such gaps, sometimes exceeding 1.0 Gb, were also identified in earlier studies with elasmobranchs (reviewed in Kuraku 2021). One possible cause may be the highly repetitive nature of the genomes. The size of the *Ch. phantasma* genome remains to be measured with non-sequence-based methods; these analyses are however hindered by the rarity of this species and the difficulty of securing live cells for karyotyping and genome size measurement.

### Chondrichthyan sex determination studies

The present study provides the first identification of DNA sequences of a candidate X chromosome in a holocephalan species (Figure 6). This sequence showed an approximately doubled sequence depth consistently in females. The sequence is only 500 Kb long and is therefore short for an entire chromosome in comparison with typical vertebrate chromosomes. In the chicken, which has numerous microchromosomes, all chromosomes are larger than 2 Mb in the latest telomere-to-telomere genome assembly (Huang & Li, 2023).

The putative X chromosome sequence of *Ch. phantasma* contains 61 predicted protein-coding genes, but none were orthologous to the master sex determination genes namely, *Sox3* (e.g., *Sry*), or *Dmrt1*, as well as a member of TGF-beta superfamily and its receptors, such as *Amh*, *Amhr2*, *Bmpr1b*, *Gsdf*, and *Gdf6*, identified in other vertebrates including mammals, birds, and teleost fishes (Bertho et al., 2021; Nagahama et al., 2021; Kitano et al., 2024). Within the Chondrichthyes, X chromosome sequences were reported in publication in *S. tigrinum R. typus*, and *C. plagiosum*, and those chromosomes were shown to be homologous to each other (Yamaguchi et al., 2023; Wu et al., 2024). Some genome assemblies of elasmobranchs in public sequence databases include chromosomes labelled as X or Y, but these identities have yet to be substantiated (Sendell-Price et al., 2023; Larivière et al., 2024). Our comparison of ortholog locations showed non-homology of the *Ch. phantasma* X chromosome sequence (Scaffold 41) to the *S. tigrinum* X chromosome (Scaffold 41, coincidentally with the same number). These species both display male heterogamety, but genomic comparisons show discordance in chromosomal origins. Investigations on other chondrichthyan lineages including batoids (rays and skates), as well as on more chimaeroids, are expected to elucidate which represents the ancestral pattern of sex chromosome organization and how this discrepancy arose. One possibility is that turnover of the sex chromosome or extensive rearrangements of the X chromosome occurred in the lineage leading to *Ch. phantasma*, after the split between the holocephalan and elasmobranch lineages. The present study used a female that should lack the Y chromosome, and further investigation are required to identify Y chromosome sequences that will provide clues for understanding the sex determination mechanism in chondrichthyans.

## Acknowledgments

We thank Kaori Tastumi for assistance in Hi-C library preparation, Yoshinobu Uno for discussion about the karyotype, Shiganori Suzuki, Yukinori Yamamoto and Haruna Yamamoto for animal sampling. This work was supported by NIG-JOINT (3B2022) to S. Hirase, “Strategic Research Projects” grant from ROIS (Research Organization of Information and Systems) to S. Kuraku, Grants-in-Aid for Scientific Research (22H00377) to K. Kikuchi and Grant-in-Aid from JSPS Fellows (DC1 202022659) to A. Teramura. Computations were partially performed on the NIG supercomputer at ROIS National Institute of Genetics.

## Data availability statement and benefit-sharing

The genome assembly of *Ch. phantasma* and the gene models associated to it are deposited at https://figshare.com/s/3ceecbdf3e6b95e17542 and https://figshare.com/s/08033f3e1cca0cb6d77e, respectively. These data, as well as the associated raw sequence reads, will be deposited in NCBI under BioProject PRJNA1075468 upon publication.

## Author contributions

This work was conceived by A.T. and K.K. The data was analyzed by A.T., M.K., and S.K. S.K. and A.T. wrote the manuscript with input from S.H. and K.K. All authors read and approved the final version of the manuscript.

